# The ABCF proteins in *Escherichia coli* individually alleviate nascent peptide-induced noncanonical translations

**DOI:** 10.1101/2023.10.04.560807

**Authors:** Yuhei Chadani, Eri Uemura, Kohei Yamazaki, Miku Kurihara, Hideki Taguchi

## Abstract

Organisms possess a wide variety of proteins with a diverse repertoire of amino acid sequences, and their synthesis relies on the ribosome. Empirical observations have led to the misconception that ribosomes are robust protein factories, but in reality, they have several weaknesses. For instance, ribosomes stall during the translation of the proline-rich sequences, but the translation elongation factor EF-P assists in synthesizing proteins containing the poly-proline sequences. Thus, living organisms have evolved to expand the translation capability of ribosomes through the acquisition of translation elongation factors, enabling the synthesis of diverse proteins.

In this study, we have revealed that *Escherichia coli* ATP-Binding Cassette family-F (ABCF) proteins, YheS, YbiT, EttA and Uup, individually cope with various noncanonical translations induced by nascent peptide sequences within the exit tunnel of the ribosome. The correspondence between noncanonical translations and ABCFs was YheS for the translational arrest by nascent SecM, YbiT for poly-basic sequence-dependent ribosome stalling and poly-acidic sequence-dependent intrinsic ribosome destabilization (IRD), EttA for IRD at the early stage of elongation, and Uup for poly-proline-dependent stalling. Our results suggest that the ATP hydrolysis-coupled structural rearrangement and interdomain linker sequence between the two nucleotide-binding domains play crucial roles in alleviating the noncanonical translations. Our study highlights a new aspect of ABCF proteins to reduce the potential risks that are encoded within the nascent peptide sequences.

**Significance statement:** Proteins, that constitute living organisms, exhibit a diverse range of amino acid sequences. However, it has become evident that ribosomes have difficulties in synthesizing certain amino acid sequences, including the poly-basic, poly-acidic, and poly-proline sequences. The mechanisms underlying the expression of proteins with such challenging sequences remain largely elusive. In this study, we have unveiled that translation factor ABCF proteins in *Escherichia coli* promote various kinds of problematic amino acid sequences that inhibit efficient translation. Through the actions of translation elongation factors including the ABCF proteins, the translation system acquires robustness in synthesizing a vast repertoire of amino acid sequences.

## Introduction

Numerous proteins in organisms show unique amino acid sequences that contribute to a wide range of cellular functions. This diversity in proteins is an essential driving force underlying the evolution of life. Therefore, the ribosome, responsible for protein synthesis, must synthesize an immense repertoire of amino acid sequences with high versatility. The ribosome itself has been considered very robust because of the variety of proteins expressed and functioning in the cells. However, recent advances have illuminated that the translation system is robust, but the ribosome itself is not universally capable of synthesizing all sequences with equal efficiency.

Ribosome-mediated translation elongation is believed to be controlled by mRNA sequences such as secondary structure and codon usage. In addition, decades of studies have demonstrated that the nascent polypeptide chain being synthesized by the ribosome itself also regulates translation. The nascent peptide interacts with the ribosomal exit tunnel and exerts various influences on ribosomal function. Particularly, in consecutive proline sequences, translation elongation is significantly hampered due to the inhibition of transpeptidation (1–3). However, organisms have evolved a solution to this challenge by obtaining a translation elongation factor EF-P (eukaryote: eIF5A), which rescues translation stalling at proline-rich motifs, enabling the synthesis of proteins containing such sequences (4–6). EF-P and eIF5A act on the peptidyl-tRNA within the P-site of the stalled ribosome from the vacant E-site, promoting transpeptidation and alleviating translation stalling (7, 8).

In addition to the poly-proline sequence, translation of poly-basic (positive charge) or poly-acidic (negative charge) sequences is also challenging for the ribosome. When a polypeptide chain passing through the ribosome tunnel contains an abundance of positively charged amino acids, translation elongation can be hindered due to interactions with the negatively charged tunnel structure (9, 10). Conversely, the translation of poly-acidic sequences can lead to difficulties in maintaining the intact ribosome complex conformation, resulting in stochastic premature termination termed intrinsic ribosome destabilization (IRD)(11–13). Thus, the growing nascent peptide can impact ribosomal activity through the interplay of ribosome structure and chemical properties.

On the other hand, several of the translation-inhibitory nascent peptides are harnessed to the survival strategies of the organisms. The nascent peptide of *Escherichia coli* SecM interacts with the ribosome tunnel, distorting the position of SecM peptidyl-tRNA itself (14, 15). This inhibits the peptidyl-transfer reaction, resulting in the translation “arrest”. This phenomenon promotes the expression of downstream SecA translocase, and the expressed SecA releases the ribosome arrested by SecM in its translocase activity-dependent manner (16). Other arrest peptides that function as “monitoring substrates” to sense membrane transport activity as well as SecM, are widely found among bacteria (17–19). Furthermore, SecM-like regulatory nascent peptides are also found in eukaryotes (20, 21), underscoring their efficacy in gene regulation.

While excepting for the regulatory nascent peptides, noncanonical translations such as translation stalling or IRD, are in general potential risks of protein synthesis. While EF-P acts on the translation of poly-proline sequences, it has not been shown whether other translation factors play similar roles in other problematic sequences. In recent years, a subfamily of the ATP Binding Cassette subfamily-F, known as ABCF proteins that are conserved from bacteria to eukaryotes, has been reported to rescue the ribosome trapped by antibiotics (22–26). Such antibiotics-resistant (ARE)-ABCF proteins are constituted from the two nucleotide-binding domains (NBDs), the interdomain linker sequence, and C-terminal extension (CTE). Once bound to the vacant E-site of the ribosome, similar to EF-P, the interdomain linker of ABCF proteins is inserted into the catalytic core of the ribosome. This rearranges the architecture of the peptidyl transferase center (PTC) and the tRNA within the P-site, effectively preventing the binding of drugs. Furthermore, it has been reported that EttA, one of ABCFs in *E. coli*, interacts with and stabilizes P-site tRNA, thereby promoting early-stage translation (27, 28). These imply that ABCF proteins have the potential to rearrange the catalytic core of translation, thereby rescuing ribosomes in various aberrant states. Although a variety of ABCF proteins have been identified across different organisms, ranging from *E. coli* to humans (26), their physiological functions remain elusive.

In this study, we hypothesized that ABCF proteins might exert inhibitory effects on various noncanonical translations induced by nascent peptides. From a series of biochemical analyses using the *E. coli* translation system, we found that each of the four ABCF proteins in *E. coli* specifically addresses different types of noncanonical translations, from potential risky sequences to the programmed arrest sequence, SecM. A class of ABCF proteins would increase the robustness of the translation system and contribute to the foundation of protein evolution.

## Results

### *E. coli* ABCF protein YheS can release the ribosome arrested by the SecM nascent peptide

Within the *E. coli* K-12 strain, there are four distinct ABCF proteins, each sharing less than 50% homology with one another (**Fig. 1A**). These proteins commonly consist of two NBDs and unique interdomain linker sequences (**Fig. 1B**). The C-terminal extension (CTE), found in several ARE-ABCFs, are also present in YheS and Uup.

**Figure 1.**
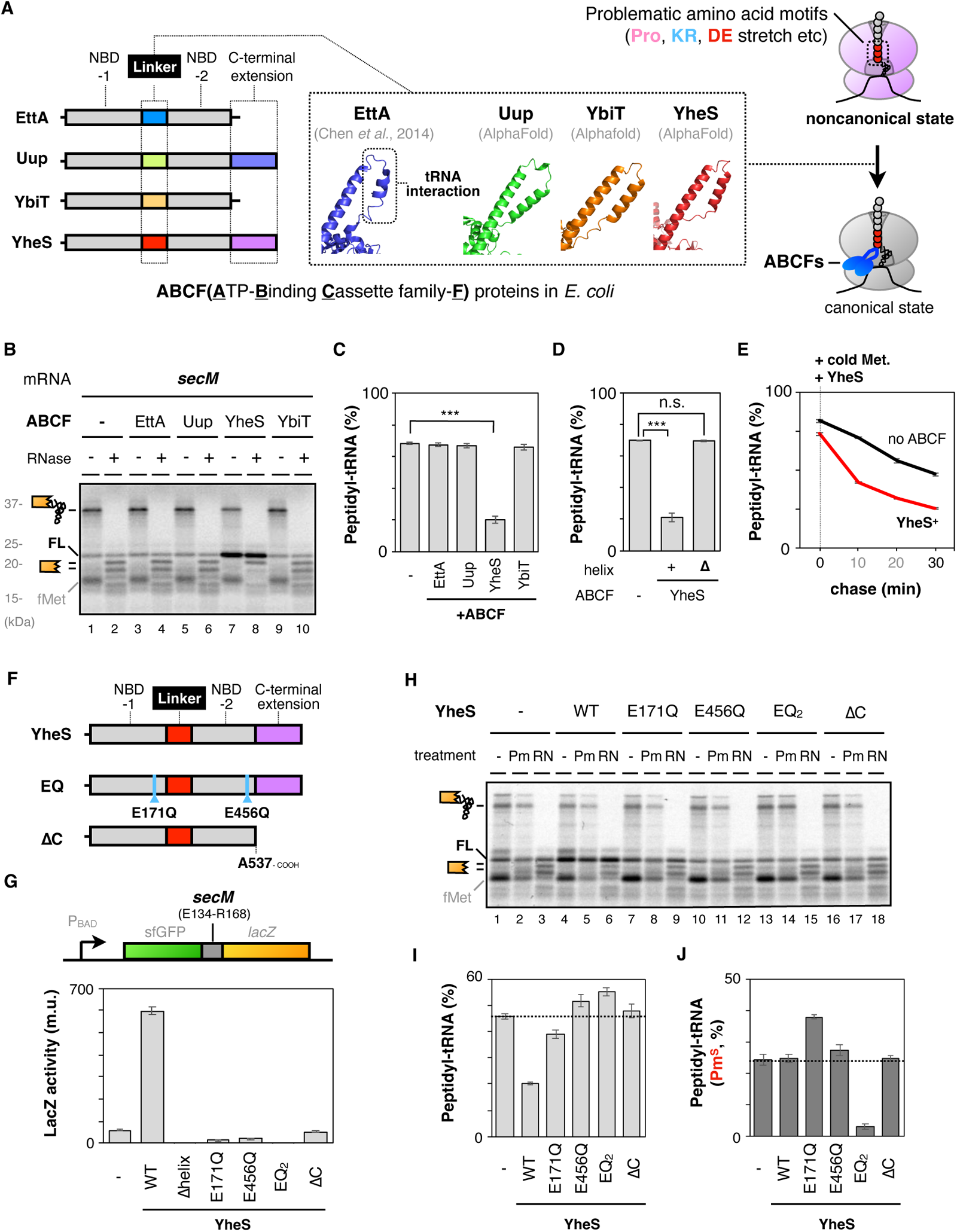
*E. coli* YheS releases the ribosome arrested by SecM nascent peptide. **A**. Four endogenous ABCF proteins, that possess characteristic interdomain linkers, individually address the ribosome in the aberrant state(s). The interdomain linker of each ABCF was described from published data (27) (PDB: 3J5S) or AlphaFold Protein Structure database (49). **B**. The *secM* mRNA was translated by PURE*frex* in the presence of Cy5-Met-tRNA and indicated ABCF proteins (final 1 µM). Samples were separated by neutral pH SDS-PAGE with optional RNase A (RN) pretreatment. The peptidyl-tRNA and tRNA-released truncated peptide are schematically indicated. “FL” stands for the full-length product. **C**. Proportion of the SecM peptidyl-tRNA quantified from gel images represented in **Fig. 1B**. ***: P-value < 0.005 (Welch’s t-test). (#) **D**. Deletion of the alpha helix abolishes the function of YheS. The proportion of the SecM peptidyl-tRNA synthesized in the presence of YheS or its derivative lacking the linker helix (**Fig. S1A**). ***: P-value < 0.005, n.s.: no significant difference (Welch’s t-test). **E**. The *secM* mRNA was translated by PURE*frex* in the presence of ^35^S-methionine for 30 min. After the addition of cold methionine (final 2 mg/ml) and purified YheS (final 1 µM), a small portion of PUREfrex mixture was withdrawn at the indicated time and mixed with an excessive amount of TCA solution. Subsequently, the ratio of the peptidyl-tRNA at the indicated time was calculated from gel images (**Fig. S1D**) and plotted. (#) **F**. Schematic of the domain structure of YheS. **G**. Arrest-releasing activity of YheS mutants. YheS or its derivatives as indicated below the graph was expressed in *E. coli* cells expressing the GFP-*secM-lacZ* reporter (schematic beyond the graph) as described in the methods. The expression level of LacZ was quantified as miller unit (m.u.). (#) **H**. The *secM* mRNA was translated by PURE*frex* in the presence of Cy5-Met-tRNA. After 10 min from the addition of YheS or its derivatives, a portion was withdrawn and mixed with the final 200 µg/ml puromycin (Pm) and further incubated for 10 min. The proportion of the SecM peptidyl-tRNA was quantified from gel images. (#). ***: P-value < 0.005 (Welch’s t-test). (#) **I**. Proportion of the SecM peptidyl-tRNA quantified from gel images represented in **Fig. 1H**. **J**. Proportion of the puromycin-reactive SecM peptidyl-tRNA quantified from gel images represented in **Fig. 1H**. (#) The mean values ±SE estimated from three independent technical replicates are shown.

Various kinds of nascent peptide sequences trigger noncanonical translations, such as ribosome stalling or premature termination. Notably, these aberrant phenomena commonly lead to the accumulation of peptidyl-tRNA molecules, detectable through gel electrophoresis analysis (11, 29). In this study, we assessed the impact of each ABCF on noncanonical translations by evaluating the accumulation of peptidyl-tRNA. Moreover, to eliminate potential indirect effects of the ABCF proteins in *E. coli*, we conducted experiments using the reconstituted *in vitro* translation system (PURE system; PURE*frex* v1.0) (30).

*E. coli* ABCF proteins exhibited effects on various noncanonical translations, which we will delve into shortly. However, what surprised us the most among the sequences we tested was the ability of YheS to release the SecM-induced translation arrest (**Fig. 1**). In the case of *secM* mRNA, most of the translation product accumulates as peptidyl-tRNA, indicating the strong arrest activity of the SecM nascent peptide (**Fig. 1B, lanes 1 and 2, Fig. 1C**). Conversely, the presence of YheS specifically enhanced the production of the full-length polypeptide (FL), indicating that ribosomes reach to the stop codon beyond the arrest peptide motif (**Fig. 1B, lanes 7 and 8**).

Previous studies have shown that the interdomain linker of ARE-ABCFs plays a critical role in rescuing the ribosomes trapped by antibiotics (23–25). Therefore, we investigated whether the interdomain linker of YheS is similarly essential for its function. Strikingly, the deletion of the α helix within the interdomain linker completely inactivated YheS (**Fig. 1D, Fig. S1A**). This result indicates that YheS releases the SecM-arrested ribosome through its interdomain linker. In addition, the arrest-releasing activity of YheS was independent of the signal sequence of SecM (**Fig. S1B and S1C**). These findings indicate that YheS employs a distinct mechanism from the well-established secretion-dependent process to release SecM-induced translation arrest (14, 16, 31). We also discovered that YheS was capable of releasing the translation arrest even after the SecM nascent peptide had already arrested the ribosome (**Fig. 1E and S1D**). In contrast, YheS had minimal impact on other ribosome-arresting peptides in *E. coli*, such as *tnaC* (32) and *speFL* (33)(**Fig. S1E and S1F**). Based on these results, we conclude that YheS efficiently rescues the ribosomes arrested by SecM, while showing limited effects on other ribosome-arresting peptides that inhibit translation through distinct mechanisms from SecM.

### Structural features required for the function of YheS

Previous studies on ARE-ABCFs have shown that several structural features were critical for their functions. For instance, the ATPase activity of NBD, which would trigger the structural rearrangements like other ABC family proteins (34–36), is indispensable for the function of ABCFs (22, 28, 37). Moreover, the C-terminal extension, also present in YheS (and Uup), is essential for the function of ARE-ABCF, VmlR (23, 37). To investigate their importance, we introduced mutations at E175 and E456 residues, which constitute the catalytic core of the ATPase activity of the two ABC cassettes, or deleted the C-terminal extension of YheS (**Fig. 1F**).

We initially employed the GFP-SecM-LacZ reporter presented in **Fig. 1G**, which expresses LacZ exclusively when YheS releases the translation arrest. The removal of the helix within the interdomain linker abolished the YheS-dependent expression of LacZ, confirming the validity of the reporter (**WT vs. Δhelix**). All E175Q, E456Q, and double mutations (EQ^2^: E175Q E456Q) completely abolished the function of YheS (**Fig. 1G**). Similarly, the absence of the C-terminal extension also eliminated the arrest-releasing activity of YheS (**Fig. 1G, ΔC**).

We further assessed the influence of these mutations using the PURE system (**Fig. 1H**). The deletion of the C-terminal extension abolished the activity of YheS, consistent with the result from the in vivo reporter assay. However, the ATPase mutants exhibited different effects from each other. The E171Q mutant retained a slight degree of arrest-releasing activity, whereas the E458Q mutant completely lost its activity (**Fig. 1I**). Intriguingly, the addition of the EQ_2_ mutant led to a more significant accumulation of peptidyl-tRNAs and nearly completely blocked the action of puromycin (**Fig. 1I and 1J**). These observations suggest that ATP hydrolysis within the two NBDs does not function symmetrically. Moreover, YheS in the initial binding state, prior to ATP hydrolysis, may not necessarily alleviate the SecM-induced translation arrest; instead, it might engage in interactions that strengthen the translation arrest. This would reflect the YheS-dependent stabilization of the SecM peptidyl-tRNA, or the distortion of peptidyl-tRNA by insertion of YheS’s interdomain linker, as observed in the studies on the ARE-ABCFs (23, 24, 26).

### EttA and YbiT alleviate intrinsic ribosome destabilization (IRD) triggered by polyacidic sequences

Given the arrest-releasing activity of YheS, we aimed to investigate whether other ABCFs also play a role in coping with nascent peptide-dependent noncanonical translations. Our previous research has demonstrated that the translation of poly-acidic sequences triggers intrinsic ribosome destabilization (IRD), leading to Pth-mediated premature termination (11, 13). For instance, over 50% of the synthesized product from a model GFP mRNA containing the 20D poly-acidic sequence accumulated as Pth-sensitive peptidyl-tRNA (**Fig. 2A, lanes 1-4, Fig. 2B**). Pth is incapable of hydrolyzing the peptidyl-tRNA within the intact 70S complex (38, 39). Therefore, the sensitivity of peptidyl-tRNA to Pth serves as a distinguishing factor between the two similar yet distinct noncanonical translations, ribosome stalling and IRD.

**Figure 2.**
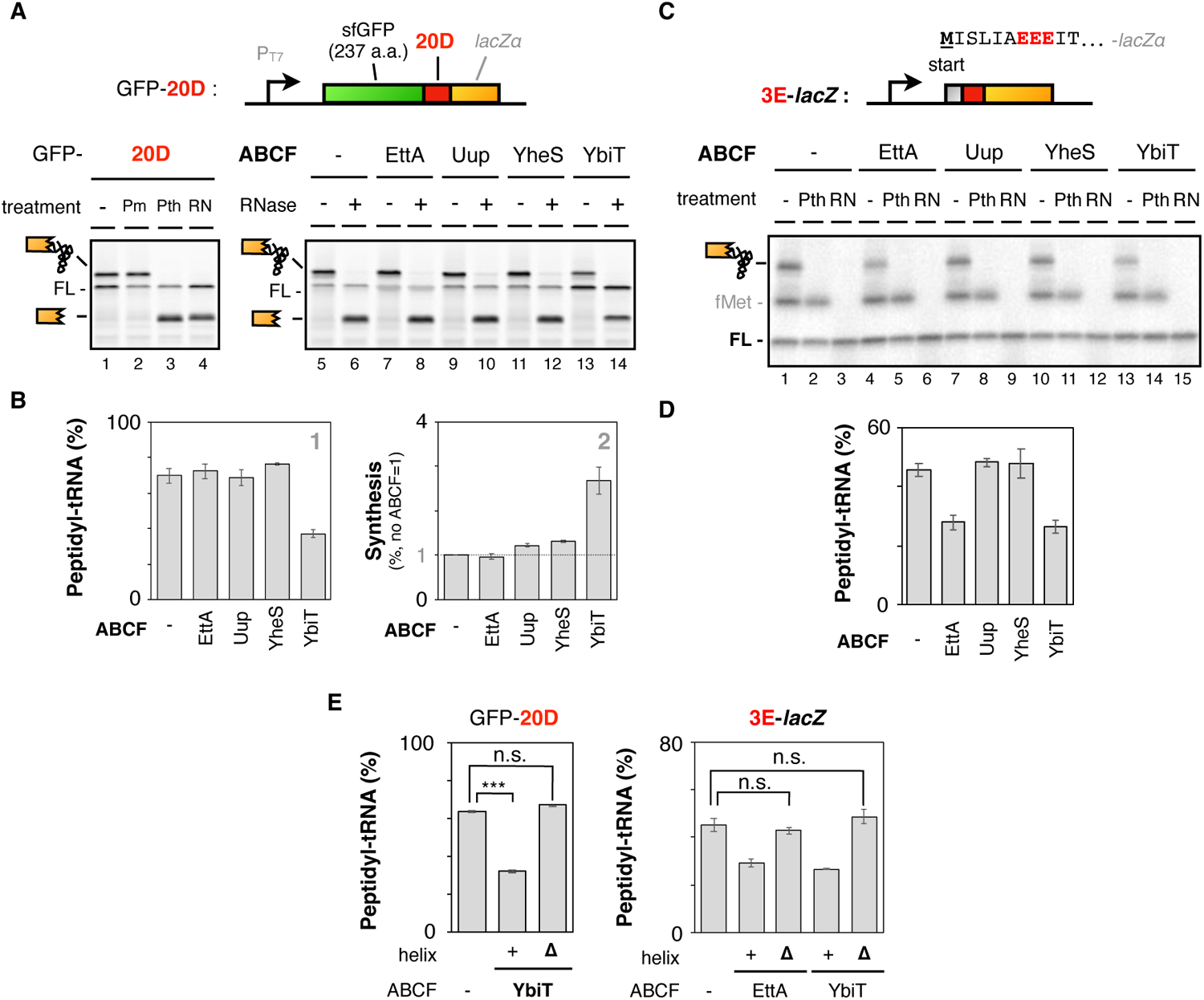
YbiT and EttA repress IRD-dependent premature termination. **A. Lanes 1-4**: The GFP-20D mRNA was translated by the PURE system (PURE*frex* v1.0) in the presence of Cy5-Met-tRNA. Samples were treated with puromycin (Pm) or peptidyl-tRNA hydrolase (Pth) as indicated, and separated by neutral pH SDS-PAGE with optional RNase A (RN) pretreatment. The peptidyl-tRNA and tRNA-released truncated peptide are schematically indicated. “FL” stands for the full-length product. **Lanes 5-14**: The GFP-20D mRNA was translated by PURE*frex* in the presence of indicated ABCF proteins (final 1 µM). **B. panel 1**: The ratio of the GFP-20D peptidyl-tRNA, calculated as described in the methods. **panel 2**: The ratio of the full-length product (FL), calculated as described in the methods. (#) **C**. The 3E-*lacZ* mRNA was translated by PURE*frex* in the presence of ^35^S-methionine and indicated ABCF proteins (final 1 µM). Samples were treated with Pth as indicated and separated by neutral pH SDS-PAGE with optional RNase A (RN) pretreatment. The peptidyl-tRNA is schematically indicated. “FL” and “fMet” stand for the full-length product and fMet-tRNA, respectively. **D**. The ratio of the 3E peptidyl-tRNA in the presence of indicated ABCF was quantified from the gel images represented in **Fig. 2C**. (#) **E**. The ratio of the GFP-20D (left panel) or 3E (right panel) peptidyl-tRNA in the presence of EttA, EttAΔhelix, YbiT, or YbiTΔhelix. n.s. : no significant difference (Welch’s t-test). (#) (#) The mean values ±SE estimated from three independent technical replicates are shown.

Consequently, GFP-20D mRNAs were subjected to translation using the PURE*frex* system, which includes purified *E. coli* ABCF proteins. Notably, the accumulation of GFP-20D peptidyl-tRNA decreased significantly upon the introduction of YbiT, one of the ABCFs, into the translation mixture (**Fig. 2A, lanes 13 and 14, Fig. 2B**). This reduction in peptidyl-tRNA accumulation correlated with an increase in the production of full-length polypeptide, indicating that YbiT enhances the ability of ribosomes to successfully navigate through the translation of the 20D sequence (**Fig. 2B, panel 2**). Similar results were observed when translating other poly-acidic 20E sequences (**Fig. S2A and S2B**) or the 56-DE repeats found in the yeast YOR054C ORF (**Fig. S2C and S2D**).

It should be noted that poly-acidic sequences like the 20D repeats examined in **Fig. 2A** are present in the eukaryotic proteome but not in the prokaryotic *E. coli* proteome. However, N-terminal DE/P-enriched sequences in *E. coli*, characterized by the presence of three or more DE/P residues within a 6 amino acid window, have been shown to induce frequent IRD (12). To assess the influence of ABCFs on the translation of N-terminal IRD sequences, we employed the 3E-*lacZ* mRNA, which encodes three consecutive glutamate residues in the N-terminal region. Once again, the inclusion of YbiT prevented IRD in the early stage of translation (**Fig. 2C, lanes 13-15, Fig. 2D**). Additionally, EttA, one of the ABCFs, exhibited an inhibitory effect on the N-terminal IRD as well (**Fig. 2C, lanes 4-6**). Identical results were obtained even when the 3E motif was substituted with the 3D motif (**Fig. S2E and S2F**). The activity of EttA and YbiT depends on the interdomain linker, as was the case for YheS (**Fig. 2E**). Intriguingly, the activity of EttA appears to be context-dependent, possibly influenced by the length of the nascent peptide within the ribosome tunnel (**Fig. S2G**). Previous reports have shown that EttA promotes the early stage of translation elongation (27, 28), which is consistent with the findings here. Collectively, these findings highlight that YbiT and EttA alleviate the poly-acidic sequence-induced IRD in a similar but distinct manner from each other.

### Uup and YbiT alleviate ribosome stalling induced by poly-proline or poly-basic sequences

Previous studies on the nascent peptide have revealed that the translation of poly-basic or poly-proline amino acid sequence hampers the translation elongation, leading to ribosome stalling (2–4, 9, 10). In this section, we investigated the influence of ABCFs on ribosome stalling using consecutive twenty lysine (20K) or ten proline (10P) sequences. The addition of YbiT promoted the translation of the 20K or 20R sequence, resulting in a twofold increase in the synthesis of full-length products (**Fig. 3A, lanes 13 and 14, Fig. 3B, panel 2, 20R in Fig. S3A and S3B**). Uup, one of the ABCFs, weakened the poly-proline dependent ribosome stalling (**Fig. 3C, lanes 9 and 10, Fig. 3D**). However, EttA and YheS did not exhibit any discernible impact on the translation of GFP-20K, 20R, and 10P mRNA sequences. The alleviation of ribosome stalling depended on the interdomain linker of Uup or YbiT, as well as other ABCFs (**Fig. S3C**). We also conducted a series of ABCF-chase experiments and found that the addition of ABCFs after the occurrence of the noncanonical translation altered the proportion of the ribosome stalling, but did not influence IRD-dependent premature termination (**Fig. S3D**). This implies that ABCFs can rescue ribosomes arrested by the nascent peptide but not those irreversibly split.

**Figure 3.**
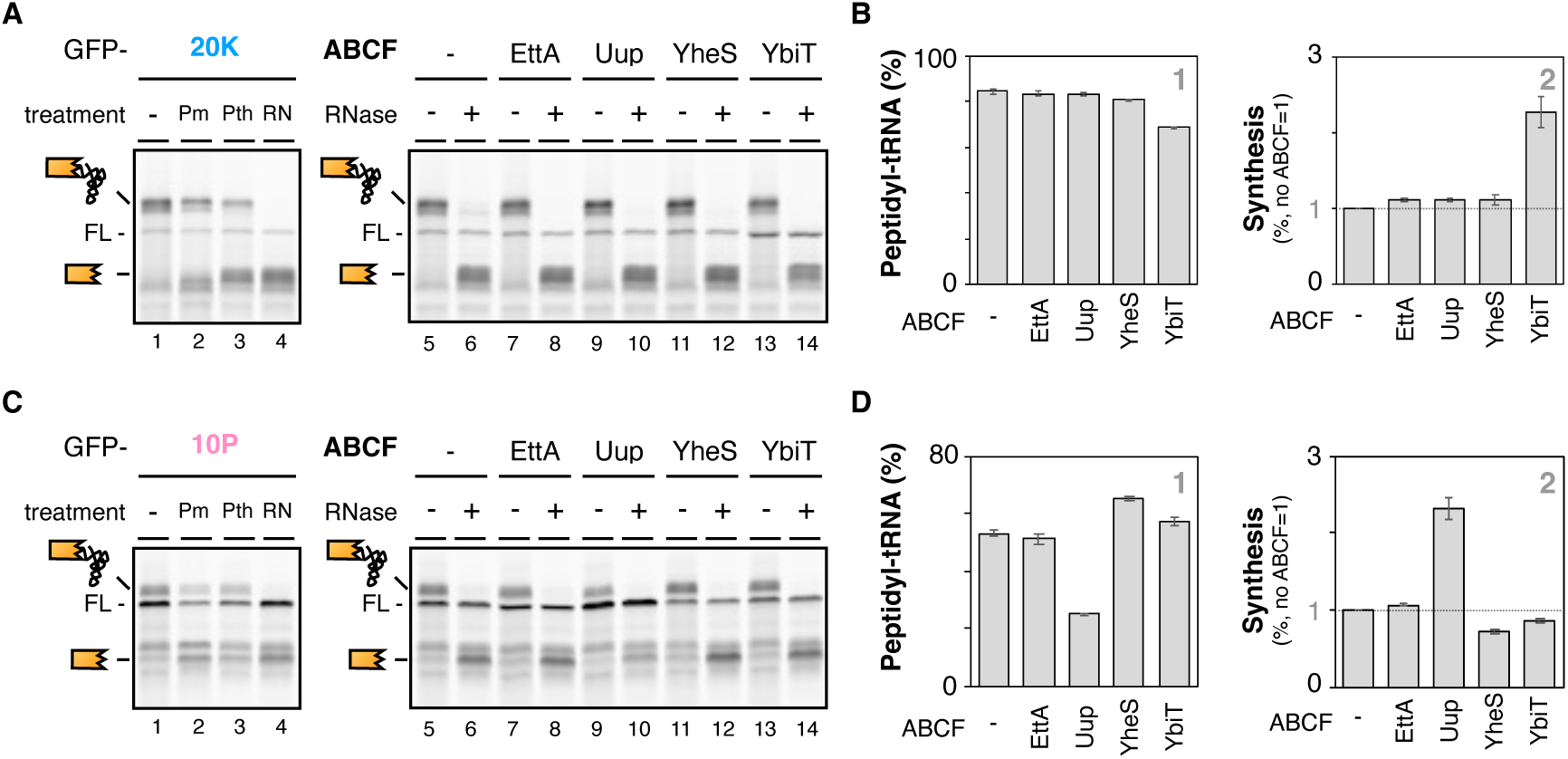
Uup and YbiT alleviate poly-proline or poly-basic sequence-dependent ribosome stalling. **A**. The translation profile of the GFP-20K mRNA, including a poly-basic amino acid sequence, was analyzed as **Fig. 2A**. **B**. Proportion of the GFP-20K peptidyl-tRNA (panel 1) and the full-length product (FL, panel 2). (#) **C**. The translation profile of the GFP-10P mRNA, including a poly-proline sequence, was analyzed as **Fig. 2A**. **E**. Proportion of the GFP-10P peptidyl-tRNA (panel 1) and the full-length product (FL, panel 2). (#) (#) The mean values ±SE estimated from three independent technical replicates are shown.

### The overall context of the interdomain linker is required for the function of YheS

The previous studies on ARE-ABCFs have shown a correlation between the characteristics of the interdomain linker and the types of antibiotics excluded from the ribosome (23, 24). Intriguingly, it has been found that the F237 residue of *Bacillus subtilis* VmlR plays a critical role in determining its specificity against antibiotics (23). *E. coli* ABCF proteins also possess unique interdomain linkers each other, and we have shown that they alleviated various noncanonical translations. Therefore, we also tested whether the amino acid residue(s) that are critical for alleviating the noncanonical translations are also present in the interdomain linker of *E. coli* ABCFs.

We employed the YheS-mediated resolution of SecM-induced translational arrest as a model. A series of alanine substitutions of the interdomain linker sequence of YheS revealed that the substitution of many residues (*e*.*g*. P291, T271-A273, and R279) attenuated the arrest-releasing activity of YheS (**Fig. 4A**). These results underscore the importance of the overall context of the interdomain linker in facilitating the release of translation arrest.

**Figure 4.**
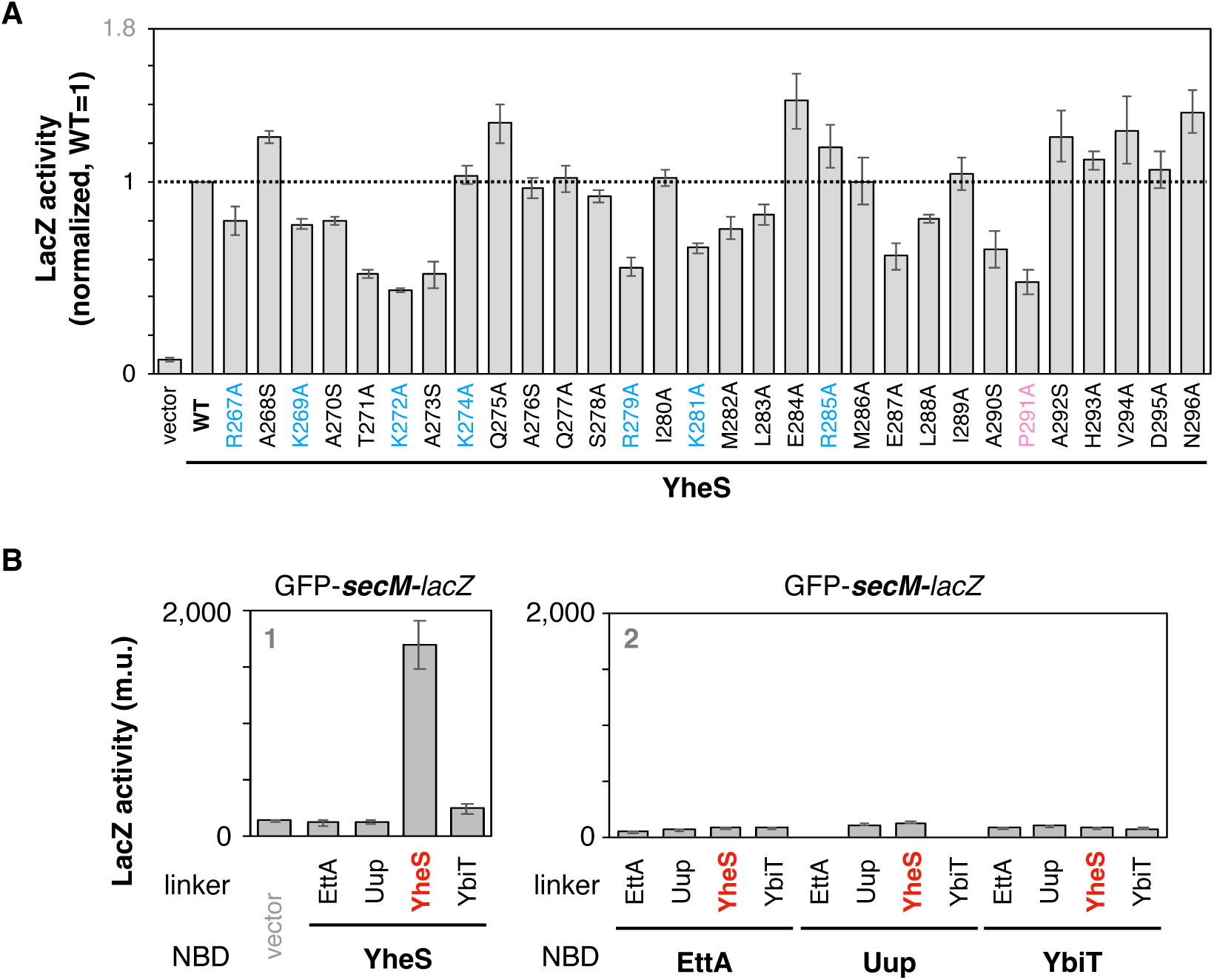
Interplay between the interdomain linker and NBDs. **A**. Alanine scanning of the YheS’s interdomain linker. The arrest-releasing activity of each YheS mutant was evaluated as shown in **Fig. 1G**. (#) **B**. The interdomain linker of each four ABCFs was comprehensively replaced by that of other ABCFs. The resulting 16 kinds of chimeric ABCFs (YheS : panel 1, others : panel 2) were individually expressed in *E. coli* cells (ECY0842, Δ*ettA*Δ*uup*Δ*yheS*Δ*ybiT*) expressing the GFP-*secM-lacZ* reporter. The expression level of LacZ was quantified as miller unit (m.u.). (#) (#) The mean values ±SE estimated from three independent biological replicates are shown.

### Interplay between the interdomain linker and NBD determines the specificity of ABCFs

Our observations have demonstrated that the interdomain linker was crucial for alleviating the noncanonical translations by ABCFs. In addition, ATP hydrolysis was also required for the arrest-releasing activity of YheS (**Fig. 1F-J**). While all four *E. coli* ABCFs possess two NBDs in common, their sequence similarity is less than 50% similarity between them. If the ATP hydrolysis-dependent structural rearrangements are indeed crucial, it is anticipated that the distinct structural features of these NBDs might influence the ability of each ABCF to alleviate noncanonical translations.

To assess this, we extensively swapped the interdomain linker among the four *E. coli* ABCFs and evaluated their function by using SecM as a model. As shown in **Fig. 4B**, replacing the interdomain linker of YheS resulted in the loss of its function, again demonstrating the importance of the interdomain linker (**panel 1**). Furthermore, swapping the interdomain linker of ABCFs other than YheS with that of YheS did not confer any arrest-releasing activity for SecM (**panel 2**). These results suggest that the arrest-releasing activity of YheS relies on the interplay between the interdomain linker and the NBDs.

## Discussion

In this study, we have successfully demonstrated that endogenous ABCF proteins in *E. coli* play a role in alleviating noncanonical translations. A key observation is that each ABCF uniquely prevents or resolves distinct noncanonical translations. From the analysis focusing on YheS, it could be inferred that the structural characteristics, including the interdomain linker that likely interacts with tRNA, the nucleotide-binding domain, and the C-terminal extension, collectively contribute to coping with noncanonical translations. Notably, the combination of the interdomain linker, which adopts unique conformations, and two NBDs define the noncanonical translations that each ABCF addresses. Furthermore, the ATP hydrolysis activity of the NBDs, which is associated with subsequent structural rearrangement of other ABC proteins (34–36), is also essential for the arrest-releasing activity of YheS. This implies that *E. coli* YheS rescues the SecM-arrested ribosome in an ATP-hydrolysis-coupled structural rearrangement, while several ARE-ABCFs are likely to dislodge the antibiotics without ATP hydrolysis (23, 24).

Structural analyses of ARE-ABCFs in Gram-negative bacteria have revealed that the differences in the interdomain linker determine what kinds of antibiotics they exclude from the ribosome. Intriguingly, the interdomain linker of ARE-ABCF induces a positional shift of the initiator tRNA, which locates within the P-site of the antibiotics-arrested ribosome (23, 24). This repositioning could explain the arrest-releasing activity of YheS observed in this study. However, one of the notable differences from previous studies is the occupation of the exit tunnel by the nascent peptide, which could constrain tRNA dynamics. Due to these constraints on structural rearrangement, ATP hydrolysis would be required for YheS to release the translation arrest induced by SecM. Further analysis is needed to determine whether ATP hydrolysis is essential for alleviating noncanonical translations or solely required for dissociation from the ribosome in the remaining three *E. coli* ABCFs.

Previous structural studies have shown that *E. coli* EttA also interacts with tRNA, restricting the dynamics of the tRNA within the P-site (27, 28). The interaction between EttA and tRNA might indeed impose constraints on the movement of the P-site tRNA, potentially leading to an inhibition of IRD. Our previous studies have shown that IRD is inhibited in situations where the dynamics of P-site tRNA is restricted by the preceding nascent chain (12, 13). Additionally, molecular simulations have indicated the possibility of P-site tRNA adopting a different positioning due to the translation of polyacidic sequence (40). Investigating the function of EttA and YbiT is an intriguing subject not only from the perspective of how these proteins promote translation but also from the standpoint of understanding the fundamental mechanism underlying IRD. However, it is important to note that neither EttA nor YbiT completely suppresses IRD, and the possibility of these proteins inhibiting IRD through secondary effects apart from their primary targets cannot be ruled out.

We have unveiled the function of the ABCFs via biochemical experiments, however, the physiological significance of endogenous ABCF proteins in *E. coli* remains unclear. Single-knockout strains for each ABCF showed almost no growth defects (**Fig. S4A**) or no alteration in the sensitivity to translation inhibitors such as erythromycin (**Fig. S4B**). However, *E. coli* strain lacking all four ABCFs exhibited a susceptibility to drugs related to translation elongation (**Fig. S4B**), suggesting that each ABCF collectively contributes to maintaining the robustness of translation elongation. Notably, previous reports have indicated the importance of EttA for long-term cultivation (28). It is anticipated that the ABCF proteins cooperatively act on ribosomes experiencing various difficulties under stress conditions. We predict that under low-stress conditions, the inherent risks associated with problematic amino acid sequences could be mitigated by other mechanisms, such as EF-P. However, under harsh circumstances, that include exposure to a specific class of antibiotics or poor nutrient availability, the tolerance threshold for these sequences might be surpassed, potentially leading to growth impairments in the absence of ABCFs. The inherent risk of noncanonical translations caused by the nascent peptide is widespread throughout the proteome, meaning that organisms certainly need to address this issue in order to survive in a variety of environments.

YheS resolves the translational arrest of SecM, which has a regulatory role in SecA expression (14, 16, 41, 42). This might appear counterintuitive, especially considering the physiological role of SecM as a monitoring substrate for the protein secretion system (43). However, an alternative perspective is that YheS could act as a rescue system in cases where the secretion-dependent arrest-releasing mechanism could not work, such as due to the loss of a signal sequence. Considering the lower expression of *yheS* mRNA (44), expression of YheS would be generally repressed not to interrupt the SecM-controlled secretion homeostasis but to be expressed in certain stress conditions. The deletion of the *yheS* gene had no influence on the SecM-induced translation arrest, consistent with this assumption (**Fig. S4C**). Further study is required to fully understand the physiological significance of YheS and other ABCFs in gene expression regulation as ARE-ABCFs (25, 45, 46).

The limited impact of ABCFs on distinct noncanonical translations suggests that each ABCF may have specialized interactions with certain structural elements or sequences of translating ribosomes. A series of analyses on YheS highlights the significance of the interplay between the interdomain linker, potentially involved in tRNA interactions, and NBDs, responsible for initiating the entire structural rearrangement, in resolving the SecM-induced translational arrest. In the future, uncovering the ATP hydrolysis-coupled movements of the interdomain linker and understanding the details of the arrest-releasing activity of YheS may make it possible to redirect the targeting of YheS towards other than SecM through re-design of the linker sequence. Such artificial modification and evolution of ABCF might enable us to manipulate genetic information encoded within the amino acid sequence, rather than within DNA.

## Materials and Methods

### *E. coli* strains, plasmids, and primers

*E. coli* strains, plasmids and oligonucleotides used in this study are listed in Tables S1, S2, and S3, respectively. The KEIO collection library (47) was obtained from the National BioResource Project (NBRP). Plasmids were constructed using standard cloning procedures and Gibson assembly. Detailed schemes are summarized in Table S2, and sequences of constructed plasmids are available in the Mendeley repository. Phage P1-mediated transduction was used to introduce the antibiotic-selection marker insertion mutation in *ettA* (JW4354), *uup* (JW0932), *yheS* (JW3315), or *ybiT* (JW0804) into BW25113. Subsequently, the FRT-Km^R^-FRT cassette was removed from the chromosome of the transductants using pCP20, as described (48).

### *In vitro* translation and product analysis

The coupled transcription-translation reaction was performed using PURE*frex* v1.0 (GeneFrontier) in the presence of ^35^S-methionine or Cy5-Met-tRNA^fMet^ at 37 °C, as described previously (11). DNA templates were prepared by PCR, as summarized in Table S4. The reaction mixture was treated with 1 µM of purified Pth for 10 min at 37 °C, or 200 µg/ml of puromycin for 10 min at 37 °C where indicated. The ABCF proteins and their derivatives, purified as described previously (27, 28), were added at the final 1 µM. The reaction was stopped by the addition of TCA, washed by ice-cooled acetone, dissolved in SDS sample buffer (62.5 mM Tris-HCl, pH 6.8, 2% SDS, 10% glycerol, 50 mM DTT) that had been treated with RNAsecure (Ambion). Finally, the sample was divided into two portions, one of which was incubated with 50 µg /ml of RNase A (Promega) at 37 °C for 30 min, and separated by a WIDE Range SDS-PAGE system (Nakalai Tesque). Images were visualized and analyzed by Amersham™ Typhoon™ scanner RGB system (GE healthcare) using 635 nm excitation laser and LPR emission filter or FLA7000 image analyzer (GE healthcare).

### β-galactosidase assay

*E. coli* cells harboring both the *lacZ* reporter plasmid and ABCF-expressing plasmid were grown overnight at 37 °C in LB medium supplemented with 100 µg/ml ampicillin and 20 µg/ml chloramphenicol. On the next day, they were inoculated into fresh LB medium containing 2x 10^-3^ % arabinose, 100 µg/ml ampicillin, and 20 µg/ml chloramphenicol and were grown at 37 °C for ∼2 hrs (A_660_ = ∼0.2). Afterward, a final 100 µM IPTG was added to express the ABCF proteins. After further inoculation for 1 hour (A_660_ = ∼0.6), 20 µl portions were subjected to a β-galactosidase assay as described previously (11, 12).

### Quantification of the signals from gel images

The ratio of the translation-completed chain (full-length: FL) against RNase-sensitive polypeptidyl-tRNAs (pep-tRNAs), which signifies the frequency of the noncanonical translation events including pausing and premature termination, was calculated by the following formula.

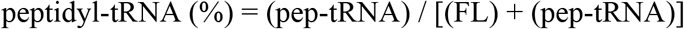

The synthesis ratio of the “Full-length” product was calculated as the ratio of the full-length product (FL) against the polypeptide part of the peptidyl-tRNAs (truncate) by the following formula. Subsequently, these values were normalized to the control experiment.

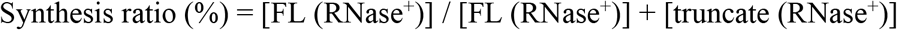

The peptidyl-tRNA that is sensitive to puromycin was then calculated with the following formula.

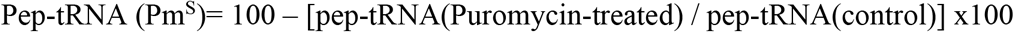

The radioactivity (^35^S-methionine) or Cy5-fluorescence proportion of “FL” and “pep-tRNA” among the samples was quantified by the Multi Gauge software (Fujifilm).

### Data analyses

Statistical analyses were conducted by using the software R (https://www.r-project.org).

### Description of the structural data

Published or predicted structural data were obtained from PDBJ (https://pdbj.org/, 3J5S) (27), or AlphaFold Protein Structure Database (https://alphafold.ebi.ac.uk, P43672, P63389, P0A9U3), respectively (49). The three-dimensional representation was generated by using PyMOL software (2.5.0).

## Data Availability

All data generated or analyzed during this study are included in the manuscript and supporting information. Raw data files are available in the Mendeley Data repository (DOI: 10.17632/x347y36n49.1).

## Acknowledgement

We would like to express our gratitude to Hikaru Takada for valuable discussions. We also thank for the Bio-support Center at Tokyo Tech for DNA sequencing. This work was supported by MEXT Grants-in-Aid for Scientific Research (Grant Numbers JP20H05925 to HT, 23H02410 to YC) and a grant from the Ohsumi Frontier Science Foundation, and a grant from the Japan Foundation for Applied Enzymology to YC.

## References

1. P. Huter, et al., Structural Basis for Polyproline-Mediated Ribosome Stalling and Rescue by the Translation Elongation Factor EF-P. Mol. Cell 68, 515–527.e6 (2017).

2. S. Ude, et al., Translation Elongation Factor EF-P Alleviates Ribosome Stalling at Polyproline Stretches. Science 339, 82–85 (2013).

3. L. K. Doerfel, et al., EF-P Is Essential for Rapid Synthesis of Proteins Containing Consecutive Proline Residues. Science 339, 85–88 (2013).

4. E. Gutierrez, et al., eIF5A Promotes Translation of Polyproline Motifs. Mol. Cell 51, 35–45 (2013).

5. A. P. Schuller, C. C.-C. Wu, T. E. Dever, A. R. Buskirk, R. Green, eIF5A Functions Globally in Translation Elongation and Termination. Mol. Cell 66, 194–205.e5 (2017).

6. V. Pelechano, P. Alepuz, eIF5A facilitates translation termination globally and promotes the elongation of many non polyproline-specific tripeptide sequences. Nucleic Acids Res. 45, 7326–7338 (2017).

7. G. Blaha, R. E. Stanley, T. A. Steitz, Formation of the First Peptide Bond: The Structure of EF-P Bound to the 70S Ribosome. Science 325, 966–970 (2009).

8. C. Schmidt, et al., Structure of the hypusinylated eukaryotic translation factor eIF-5A bound to the ribosome. Nucleic Acids Res. 44, 1944–1951 (2016).

9. L. N. Dimitrova, K. Kuroha, T. Tatematsu, T. Inada, Nascent Peptide-dependent Translation Arrest Leads to Not4p-mediated Protein Degradation by the Proteasome. J. Biol. Chem. 284, 10343–10352 (2009).

10. J. Lu, C. Deutsch, Electrostatics in the Ribosomal Tunnel Modulate Chain Elongation Rates. J. Mol. Biol. 384, 73–86 (2008).

11. Y. Chadani, et al., Intrinsic Ribosome Destabilization Underlies Translation and Provides an Organism with a Strategy of Environmental Sensing. Mol. Cell 68, 528–539.e5 (2017).

12. Y. Chadani, et al., Nascent polypeptide within the exit tunnel stabilizes the ribosome to counteract risky translation. EMBO J. e108299 (2021).

13. Y. Ito, et al., Nascent peptide-induced translation discontinuation in eukaryotes impacts biased amino acid usage in proteomes. Nat. Commun. 13, 7451 (2022).

14. H. Nakatogawa, K. Ito, The Ribosomal Exit Tunnel Functions as a Discriminating Gate. Cell 108, 629–636 (2002).

15. S. Bhushan, et al., SecM-Stalled Ribosomes Adopt an Altered Geometry at the Peptidyl Transferase Center. PLoS Biol. 9, e1000581 (2011).

16. H. Nakatogawa, K. Ito, Secretion Monitor, SecM, Undergoes Self-Translation Arrest in the Cytosol. Mol. Cell 7, 185–192 (2001).

17. E. Ishii, et al., Nascent chain-monitored remodeling of the Sec machinery for salinity adaptation of marine bacteria. Proc. Natl. Acad. Sci. USA 112, E5513–E5522 (2015).

18. K. Sakiyama, N. Shimokawa-Chiba, K. Fujiwara, S. Chiba, Search for translation arrest peptides encoded upstream of genes for components of protein localization pathways. Nucleic Acids Res. 49, gkab024.(2021).

19. S. Chiba, A. Lamsa, K. Pogliano, A ribosome-nascent chain sensor of membrane protein biogenesis in Bacillus subtilis. EMBO J. 28, 3461–3475 (2009).

20. H. Onouchi, et al., Nascent peptide-mediated translation elongation arrest coupled with mRNA degradation in the CGS1 gene of Arabidopsis. Genes Dev. 19, 1799–1810 (2005).

21. K. Yanagitani, Y. Kimata, H. Kadokura, K. Kohno, Translational Pausing Ensures Membrane Targeting and Cytoplasmic Splicing of XBP1u mRNA. Science 331, 586–589 (2011).

22. V. Murina, M. Kasari, V. Hauryliuk, G. C. Atkinson, Antibiotic resistance ABCF proteins reset the peptidyl transferase centre of the ribosome to counter translational arrest. Nucleic Acids Res. 46, gky050. (2018).

23. C. Crowe-McAuliffe, et al., Structural basis for antibiotic resistance mediated by the Bacillus subtilis ABCF ATPase VmlR. Proc. Natl. Acad. Sci. USA. 115, 8978–8983 (2018).

24. C. Crowe-McAuliffe, et al., Structural basis of ABCF-mediated resistance to pleuromutilin, lincosamide, and streptogramin A antibiotics in Gram-positive pathogens. Nat. Commun. 12, 3577 (2021).

25. N. Obana, et al., Genome-encoded ABCF factors implicated in intrinsic antibiotic resistance in Gram-positive bacteria: VmlR2, Ard1 and CplR. Nucleic Acids Res. 51, 4536–4554 (2023).

26. V. Murina, et al., ABCF ATPases Involved in Protein Synthesis, Ribosome Assembly and Antibiotic Resistance: Structural and Functional Diversification across the Tree of Life. J. Mol. Biol. 431, 3568–3590 (2019).

27. B. Chen, et al., EttA regulates translation by binding the ribosomal E site and restricting ribosome-tRNA dynamics. Nat. Struct. Mol. Biol. 21, 1–11 (2014).

28. G. Boël, et al., The ABC-F protein EttA gates ribosome entry into the translation elongation cycle. Nat. Struct. Mol. Biol. 21, 143–151 (2014).

29. Y. Chadani, T. Niwa, S. Chiba, H. Taguchi, K. Ito, Integrated in vivo and in vitro nascent chain profiling reveals widespread translational pausing. Proc. Natl. Acad. Sci. USA. 113, E829–838 (2016).

30. Y. Shimizu, et al., Cell-free translation reconstituted with purified components. Nat. Biotechnol. 19, 751–755 (2001).

31. M. E. Butkus, L. B. Prundeanu, D. B. Oliver, Translocon “pulling” of nascent SecM controls the duration of its translational pause and secretion-responsive secA regulation. J. Bacteriol. 185, 6719–6722 (2003).

32. F. Gong, C. Yanofsky, Instruction of Translating Ribosome by Nascent Peptide. Science 297, 1864–1867 (2002).

33. A. H. del Valle, et al., Ornithine capture by a translating ribosome controls bacterial polyamine synthesis. Nat. Microbiol. 5, 554–561 (2020).

34. D. Barthelme, et al., Ribosome recycling depends on a mechanistic link between the FeS cluster domain and a conformational switch of the twin-ATPase ABCE1. Proc. Natl. Acad. Sci. USA. 108, 3228–3233 (2011).

35. T. Becker, et al., Structural basis of highly conserved ribosome recycling in eukaryotes and archaea. Nature 482, 501–506 (2012).

36. P. Vergani, S. W. Lockless, A. C. Nairn, D. C. Gadsby, CFTR channel opening by ATP-driven tight dimerization of its nucleotide-binding domains. Nature 433, 876–880 (2005).

37. E. Jacquet, et al., ATP Hydrolysis and Pristinamycin IIA Inhibition of the Staphylococcus aureus Vga(A), a Dual ABC Protein Involved in Streptogramin A Resistance. J. Biol. Chem. 283, 25332–25339 (2008).

38. N. de Groot, A. Panet, Y. Lapidot, Enzymatic hydrolysis of peptidyl-tRNA. Biochem. Biophys. Res. Commun. 31, 37–42 (1968).

39. J. R. Menninger, M. C. Mulholland, W. S. Stirewalt, Peptidyl-tRNA hydrolase and protein chain termination. Biochim. Biophys. Acta. 217, 496–511 (1970).

40. S. E. Leininger, et al., Ribosome Elongation Kinetics of Consecutively Charged Residues Are Coupled to Electrostatic Force. Biochemistry-us 60, 3223–3235 (2021).

41. A. Murakami, H. Nakatogawa, K. Ito, Translation arrest of SecM is essential for the basal and regulated expression of SecA. Proc. Natl. Acad. Sci. USA. 101, 12330–12335 (2004).

42. H. Nakatogawa, A. Murakami, H. Mori, K. Ito, SecM facilitates translocase function of SecA by localizing its biosynthesis. Genes Dev. 19, 436–444 (2005).

43. K. Ito, H. Mori, S. Chiba, Monitoring substrate enables real-time regulation of a protein localization pathway. FEMS Microbiol. Lett. 365, fny109 (2018).

44. G.-W. Li, D. Burkhardt, C. Gross, J. S. Weissman, Quantifying Absolute Protein Synthesis Rates Reveals Principles Underlying Allocation of Cellular Resources. Cell 157, 624–635 (2014).

45. H. Takada, et al., Expression of Bacillus subtilis ABCF antibiotic resistance factor VmlR is regulated by RNA polymerase pausing, transcription attenuation, translation attenuation and (p)ppGpp. Nucleic Acids Res. 50, 6174–6189 (2022).

46. C. R. Fostier, et al., Regulation of the macrolide resistance ABC-F translation factor MsrD. Nat. Commun. 14, 3891 (2023).

47. T. Baba, et al., Construction of Escherichia coli K-12 in-frame, single-gene knockout mutants: the Keio collection. Mol Syst Biol 2, 2006.0008 (2006).

48. K. A. Datsenko, B. L. Wanner, One-step inactivation of chromosomal genes in Escherichia coli K-12 using PCR products. Proc. Natl. Acad. Sci. USA. 97, 6640–6645 (2000).

49. M. Varadi, et al., AlphaFold Protein Structure Database: massively expanding the structural coverage of protein-sequence space with high-accuracy models. Nucleic Acids Res. 50, D439–D444 (2021).

