## Supplementary Fig. S1-S4 for "The ABCF proteins in *Escherichia coli* individually alleviate nascent peptide-induced noncanonical translations"

**This PDF file includes Figures S1-S4.**

**Other supporting material for this manuscript includes Tables S1-S4.**

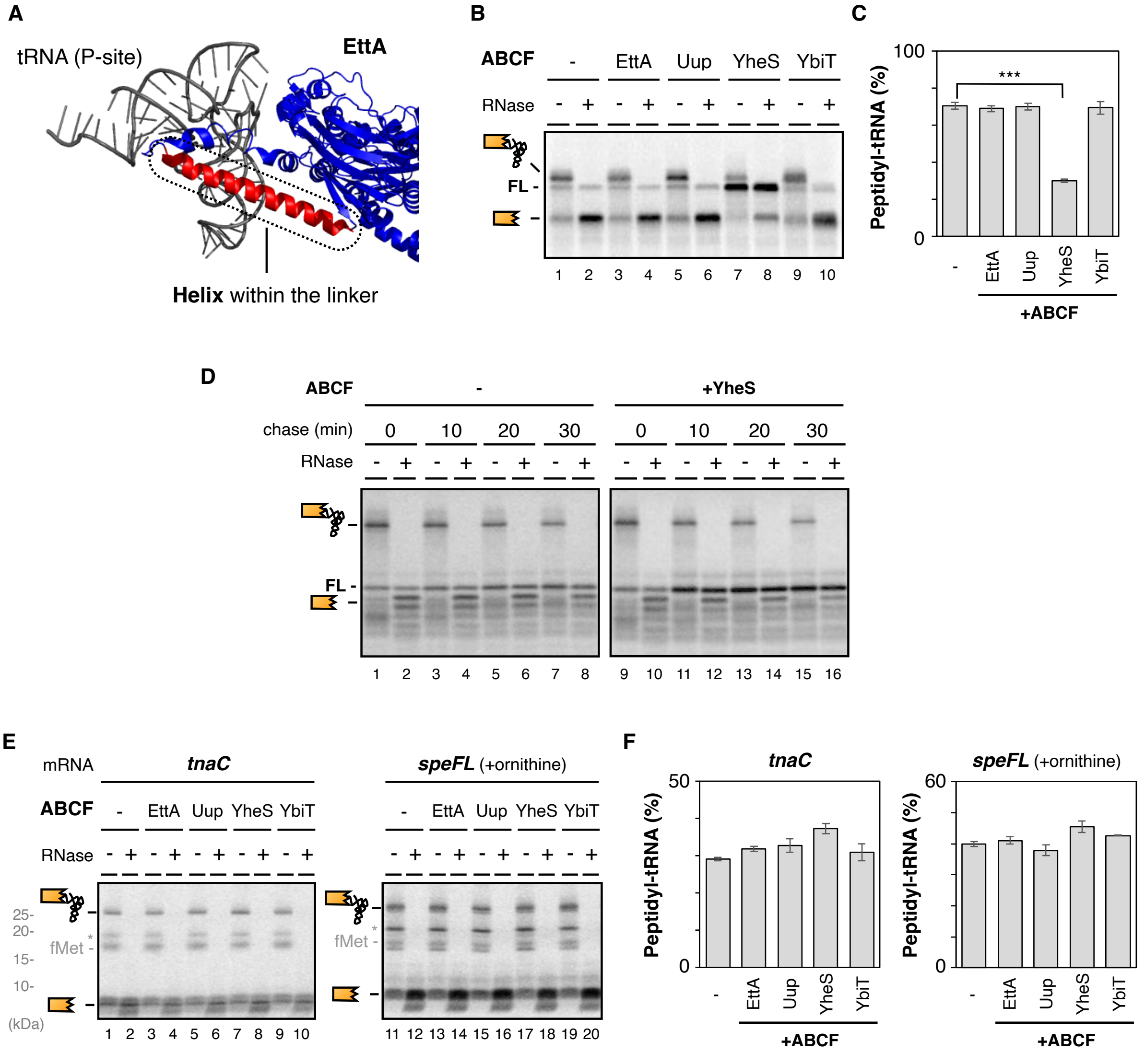


**Figure S1.**

**A**. Structural description of the interdomain linker of EttA, interacting with the P-site tRNA (27)(PDB: 3J5S). The alpha helix deleted in the following experiment is indicated by red color.

**B**. The GFP-*secM* mRNA, which lacks the N-terminal signal sequence of SecM, was translated by PURE*frex* in the presence of Cy5-Met-tRNA and indicated ABCF proteins (final 1 μM). Samples were separated by neutral pH SDS-PAGE with optional RNase A (RN) pretreatment. The peptidyl-tRNA and tRNA-released truncated peptide are schematically indicated. "FL" stands for the full-length product.

**C**. Proportion of the GFP-SecM peptidyl-tRNA quantified from gel images represented in **Fig. S1B**. ***: P-value < 0.005 (Welch's t-test). (#)

**D**. Representative gel image of YheS-chase experiments shown in **Fig. 1E**.

**E**. The *tnaC* (lanes 1-10) or *speFL* mRNA (lanes 11-20, in the presence of 10 mM ornithine) was translated by PURE*frex* in the presence of $^{35}$S-methionine and indicated ABCF proteins (final 1 μM).

**F**. Proportion of the TnaC (left) or SpeFL peptidyl-tRNA (right) quantified from gel images represented in **Fig. S1E**. (#)

(#) The mean values ±SE estimated from three independent biological or technical replicates are shown.

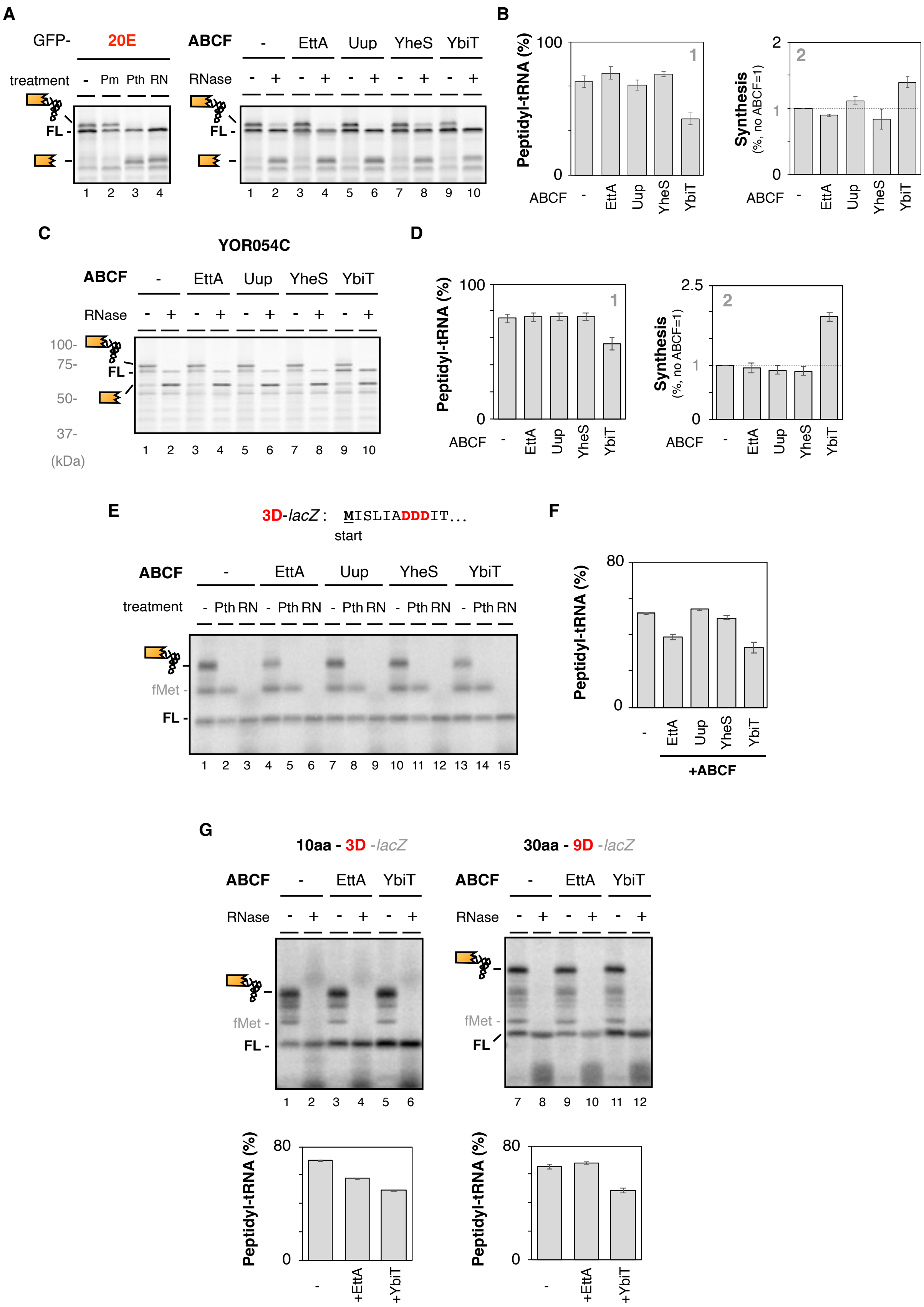


**Figure S2.**

**A**. The translation profile of the GFP-20E mRNA was analyzed as **Fig. 2A**. (#)

**B**. Proportion of the GFP-20E peptidyl-tRNA (panel 1) and the full-length product (FL, panel 2) (#).

**C**. Translation profile of the yeast YOR054C mRNA, which includes the 56-DE repeats (606th-DDDDDDDDDDDDDDDDDDDDDDDDDDDDDDDDDDDDDDDDDDDDDDDDDEDDEDEDEDDE - 661st) was analyzed as **Fig. 2A**.

**D**. Proportion of the 56-DE peptidyl-tRNA (panel 1) and the full-length product (FL, panel 2) (#).

**E**. The 3D-*lacZ* mRNA was translated by PURE*frex* in the presence of $^{35}$S-methionine and indicated ABCF and its derivatives (final 1 μM). Samples were treated with Pth as indicated, and separated by neutral pH SDS-PAGE with optional RNase A (RN) pretreatment. The peptidyl-tRNA is schematically indicated. "FL" and "fMet" stand for full-length product and fMet-tRNA, respectively.

**F**. The ratio of the 3D peptidyl-tRNA in the presence of indicated ABCF was quantified from the gel images represented in **Fig. S2E**. (#)

**G**. **Upper**: The 10aa-3D-*lacZ* (short nascent peptide within the tunnel) or 30aa-9D-*lacZ* mRNA (long nascent peptide within the tunnel) was translated by PURE*frex* in the presence of $^{35}$S-methionine and indicated ABCF (final 1 μM). **Lower**: The ratio of the 3D or 9D peptidyl-tRNA in the presence of indicated ABCF quantified from the gel images. (#)

(#) The mean values ±SE estimated from three independent technical replicates are shown.

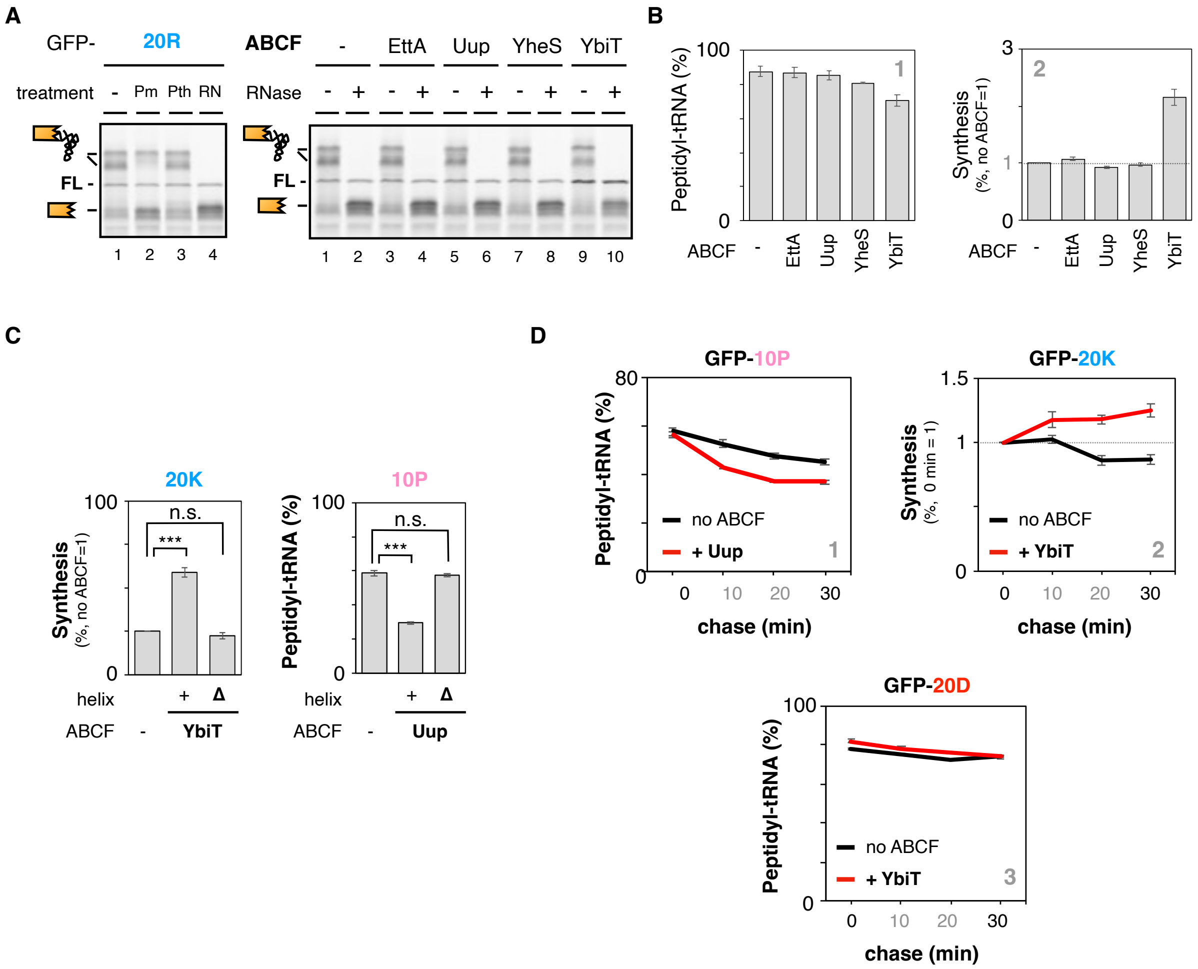


**Figure S3.**

**A**. The translation profile of the GFP-20R mRNA was analyzed as **Fig. 2A**.

**B**. Proportion of the GFP-20R peptidyl-tRNA (panel 1) and the full-length product (FL, panel 2) quantified from gel images represented in **Fig. S3A** (#).

**C**. The GFP-20K (left), or GFP-10P mRNA (right) was translated by PURE*frex* in the presence of the ABCF with or without the linker helix. The ratio of peptidyl-tRNA (10P) or that of the full-length product (20K) was calculated and plotted. ***: P-value < 0.005, n.s.: no significant difference (Welch's t-test). (#)

**D**. Chase experiment. The GFP-20D (panel 1), GFP-10P (panel 2), or GFP-20K mRNA (panel 3) was translated by PURE*frex* in the presence of $^{35}$S-methionine for 30 min. After the addition of cold methionine (final 2 mg/ml) and purified ABCF (final 1 µM), a small portion of the PURE*frex* mixture was withdrawn at the indicated time and mixed with an excessive amount of TCA solution. Then the ratio of the peptidyl-tRNA (20D, 10P) or that of the full-length product (20K) was calculated and plotted (#).

(#) The mean values ±SE estimated from three independent technical replicates are shown.

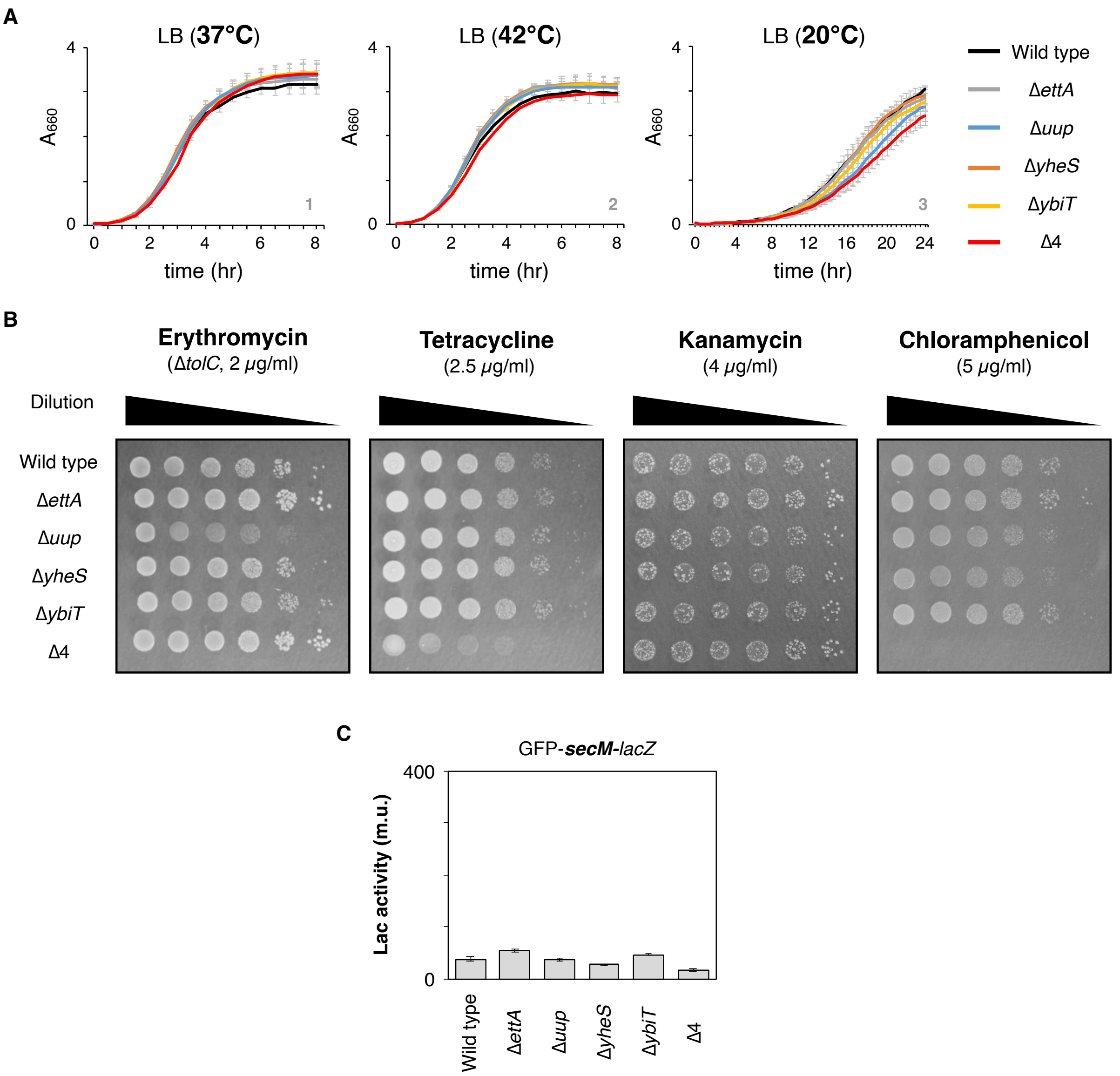


**Figure S4.**

**A**. Growth of *E. coli* strains lacking each ABCF or all of them. *E. coli* cells grown in LB medium overnight at 37 °C were diluted into fresh LB, and further cultivated at the indicated temperature with shaking (180 rpm). The $A_{660}$ values were monitored at the indicated time. (#)

**B**. Contribution of endogenous ABCFs on antibiotic resistance. Serially diluted *E. coli* cells were spotted on LB agar plates containing the indicated concentrations of the antibiotics. Note that Δ*tolC* variants were used in the case of the assessment for erythromycin resistance. Representatives of three independent experiments were shown.

**C**. The GFP-*secM-lacZ* reporter was expressed in *E. coli* cells indicated below the graph, and the expression level of LacZ was quantified as miller unit (m.u.). The Δ4 strain (ECY0842, Δ*ettA*Δ*uup*Δ*yheS*Δ*ybiT*) lacks all four ABCF genes. (#)

(#) The mean values ±SE estimated from three independent biological replicates are shown.
